# Small RNAs reflect grandparental environments in apomictic dandelion

**DOI:** 10.1101/099572

**Authors:** Lionel Morgado, Veronica Preite, Carla Oplaat, Sarit Anava, Julie Ferreira de Carvalho, Oded Rechavi, Frank Johannes, Koen J.F. Verhoeven

## Abstract

Plants can show long-term effects of environmental stresses and in some cases a stress ‘memory’ has been reported to persist across generations, potentially mediated by epigenetic mechanisms. However, few documented cases exist of transgenerational effects that persist for multiple generations and it remains unclear if or how epigenetic mechanisms are involved. Here we show that the composition of small regulatory RNAs in apomictic dandelion lineages reveals a footprint of drought stress and salicylic acid treatment experienced two generations ago. Overall proportions of 21nt and 24nt RNA pools were shifted due to grandparental treatments. While individual genes did not show strong up- or downregulation of associated sRNAs, the subset of genes that showed the strongest shifts in sRNA abundance was significantly enriched for several GO terms including stress-specific functions. This indicates that a stress-induced signal was transmitted across multiple unexposed generations leading to persistent and functional changes in epigenetic gene regulation.

---

Stress exposure triggers responses that are mediated by changes in gene regulation (Cramer et al. 2011; Heil 2002; Shao et al. 2008). In plants, some responses to environmental stresses are long-lived. For instance, upon mild pathogen infection, plants can enter a “primed” state which is expressed as a quicker or more vigorous defense response upon a second infection later in life (Conrath et al. 2002). Defense-related priming effects, and also responses to other environmental triggers, have been demonstrated to persist into offspring generations in some cases (Agrawal 2002; Mandal et al. 2012; Slaughter et al. 2012; Wang et al. 2016).

Although several different mechanisms can underlie inherited environmental effects in plants (Crisp et al. 2016), epigenetic mechanisms are considered prime candidates because of their potential for environmental sensitivity (Dowen et al. 2012) and transgenerational stability (Cortijo et al. 2014). Especially DNA methylation can be transgenerationally stable in plants and this mechanism is often proposed to mediate environmental effects that persist for multiple generations (Bilichak et al. 2015; Boyko et al. 2010; Boyko et al. 2007; Ou et al. 2012; Verhoeven et al. 2010) although empirical support for this hypothesis remains scarce (Pecinka and Scheid 2012).

Accumulating evidence indicates that regulatory small RNAs (sRNAs) also have a role in plant transgenerational effects. Indeed, sRNA biogenesis mutants in *A. thaliana* show compromised transgenerational herbivore resistance (Rasmann et al. 2012), suggesting that sRNAs are required to sustain priming responses across generations. Changes in sRNA composition have also been demonstrated in a number of species in response to heat (Bilichak et al. 2015; Ito et al. 2011; Song et al. 2016), drought (Matsui et al. 2008; Tricker et al. 2012), salinity (Borsani et al. 2005; Ding et al. 2009; Matsui et al. 2008; Song et al. 2016), cold and osmotic stress (Song et al. 2016). In some cases, these sRNA alternations have been shown to persist in the offspring of stressed plants. The mechanisms that maintain changes of sRNAs across generations remain largely unclear but may involve feed-forward interactions with (transiently) heritable DNA methylation changes (Wibowo et al. 2016).

Here, we used apomictic dandelions (*Taraxacum officinale*) to test the impact of environmental stress on sRNA composition in unexposed offspring two generations after stress treatment. Due to apomictic reproduction dandelion offspring are considered clonal copies (Bicknell and Koltunow 2004) allowing for multi-generation experiments without confounding effects of genetic differences between samples. In this species, first-generation offspring of stress-exposed plants have previously been demonstrated to show modified phenotypes and DNA methylation patterns, suggesting potential for environment-induced transgenerational epigenetic inheritance (Verhoeven et al. 2010; Verhoeven and van Gurp 2012).

We grew first-generation (G1) plants under either drought stress, salicylic acid exposure (SA; a plant hormone that is involved in several processes including defense signaling in response to pathogens; Vicente and Plasencia 2011), or under control conditions. Second (G2) and third (G3) generation apomictic progenies were obtained by single-seed descent for four replicate lineages per experimental group and were grown under common control conditions (Supplementary text: **S1**). sRNAs were sequenced at generation G3 in four individual plants per experimental group (Supplementary text: **S1-S2**). As no reference assembly currently exists for dandelion, we first assessed differences in sRNA composition between treatment groups and control using the total sRNA libraries. We found significant shifts in all size classes of sRNAs ranging from 21 to 24 nt (**Fig. 1A** and **1D**, Supplementary text: **S3**). The most pronounced changes occurred for sRNAs of size 24nt whose relative abundance in the total sRNA population was markedly reduced in both of the stress conditions compared to the control. Relative loss of TE-associated 24nt sRNA has been reported for a variety of biotic and abiotic stressors (Dowen et al. 2012; Lunardon et al. 2016; McCue et al. 2012). These changes appear to be mainly the result of hypomethylation and loss of RdDM targeting of transposable element (TE) sequences (McCue et al. 2013; Tran et al. 2005). Loss of 24 nt sRNAs is typically accompanied by gains in 21 nt sRNAs due to an increase in the transcription of precursors for this class of sRNA (Dowen et al. 2012; McCue et al. 2012). Consistent with this, the relative loss in 24 nt sRNAs after drought stress co-occurred with gains in 21 nt sRNAs, although a similar trend could not be observed for the SA treatment (**Fig. 2A** and **2D**).

**Figure 1.**
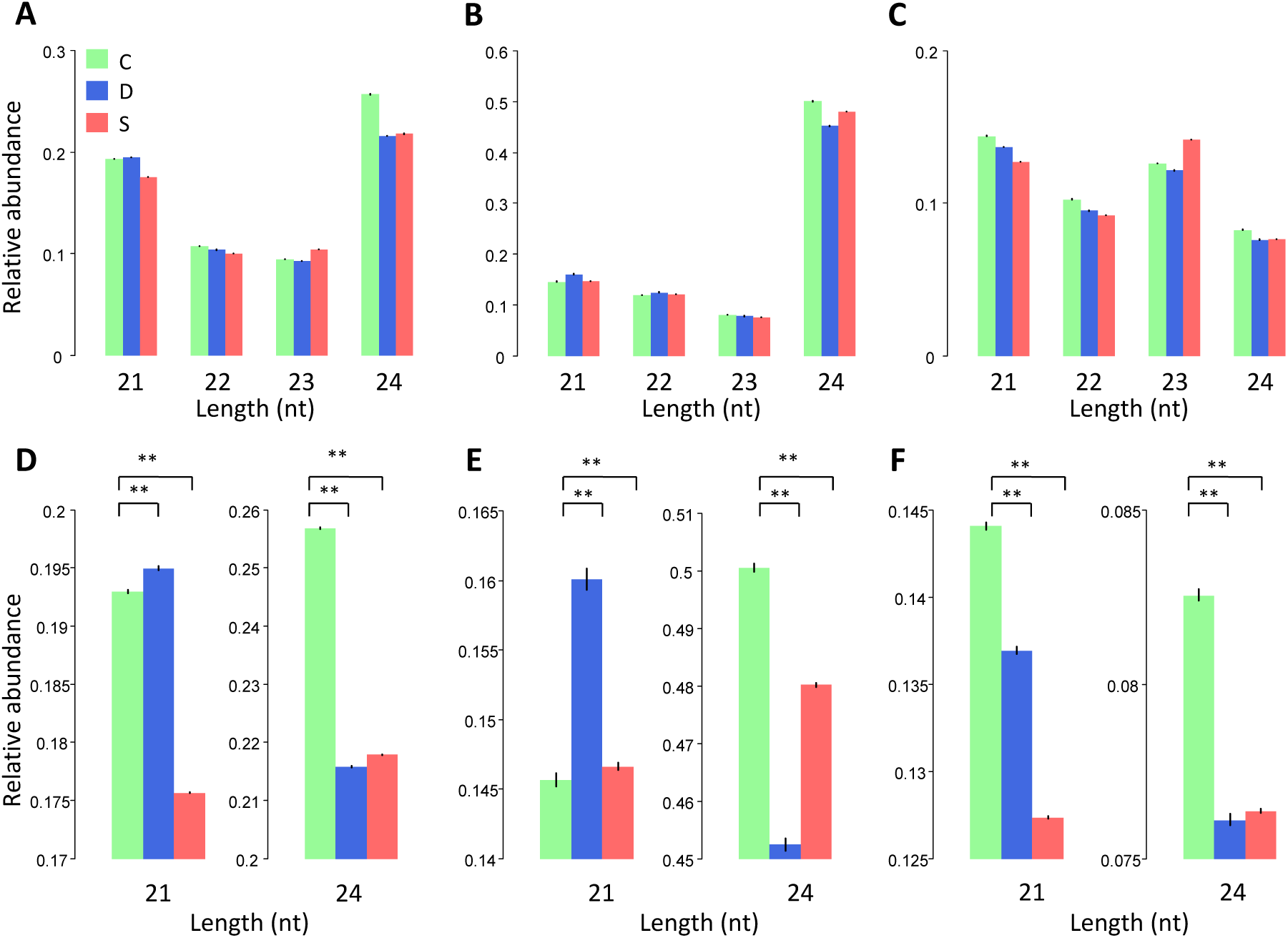
Length composition for the read libraries: all sRNAs (A and D), mapped to annotated TEs (B and E) and mapped to gene-annotated transcripts (C and F). The plots in the bottom (D, E and F) show the p-value for the bootstrap analysis performed for 21nt and 24 nt sRNA. Error bars, 95% bootstrap confidence intervals. ** p value = 10^−4^. Treatment groups (C: control; D: drought; S: salicylic acid) refer to grandparental treatments.

**Figure 2.**
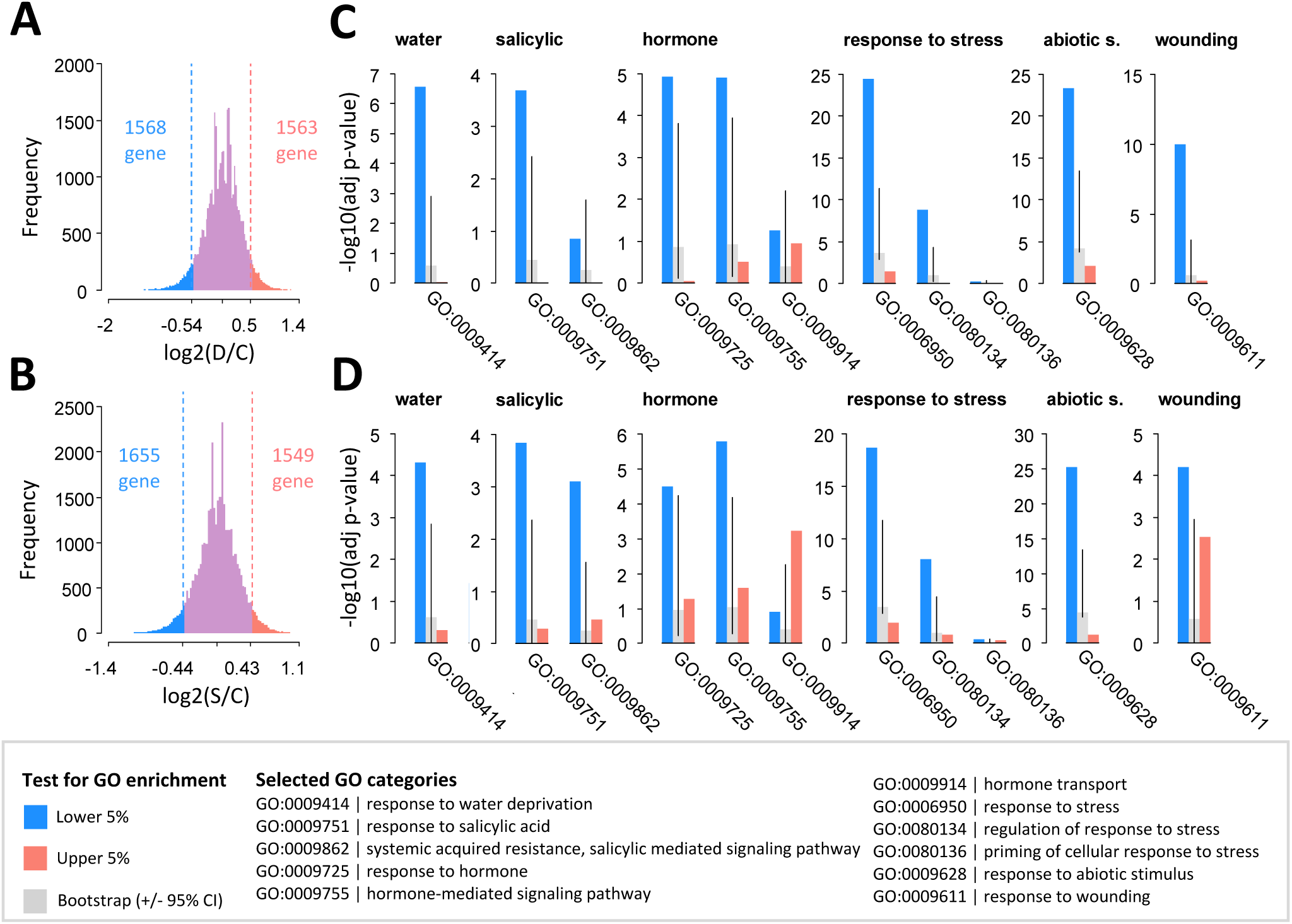
Distribution for sRNA fold change in gene-mapping transcripts after grandparental drought stress (A) and salicylic acid (B) against control. Bar plots show GO terms enrichment compared to a bootstrap analysis in the case of the drought (C) and the salicylic acid (D) sets. Error bars, 95% bootstrap confidence intervals.

The fact that the libraries show signatures of known TE-associated stress-induced changes in sRNA (at least for the drought treatment) suggests that the observed effects are dominated by changes within TE sequences. To test this more directly we took advantage of a recent TE database that was generated based on de novo clustering of repetitive sequences from the *T. officinale* genome (Ferreira de Carvalho et al. 2016a)(SI text: **S2**). We aligned sRNAs to these TE sequences, and compared the relative size abundance across conditions. A loss of 24 nt sRNAs was also observed in these TE-annotated sequences (**Fig. 2B** and **2E**). Moreover, the gain in 21 nt sRNAs was now clearer and occurred in both stress treatments, suggesting that stress-induced increases in 21 nt sRNA are subtle and may have been obscured by the complexity of the total sRNA libraries.

It is unclear how and if such sRNA shifts impact gene expression. Previous studies have reported that TEs proximal to or overlapping genes can affect transcription under stress, probably as a result of RdDM-mediated DNA methylation loss (Dowen et al. 2012; Hollister and Gaut 2009; Lister et al. 2008; Quadrana et al. 2016; Wang et al. 2013). Another mechanism by which sRNAs can affect gene expression is through *trans*-acting post-transcriptional modifications (Borsani et al. 2005). We recently assembled the complete transcriptome of dandelion (Ferreira de Carvalho et al. 2016b). Over 13500 genes could be annotated by homology with *A. thaliana.* We aligned our sRNA libraries to these transcriptomes (SI text: **S2**). On average about 53% of the reads from each library met our quality control criteria and could be successfully aligned. In contrast to TE-aligned sRNAs, the relative abundance of transcriptome-aligned sRNAs was reduced after grandparental stress exposure not only for 24nt but also for 21nt sRNAs (**Fig. 2C** and **2F**).

The observed loss of transcript-associated 24 nt sRNAs was enigmatic, as 24nt sRNA are typically depleted in genic sequences. To explore this issue in more detail, we studied the density distribution of 24nt sRNA along our annotated transcripts. Our analysis shows that 24 nt sRNAs mapped more frequently towards the 5’ and 3’ flanks of the genes, suggesting vestiges of a sRNA signal that originates from sequences outside of gene bodies, such as promoters or intergenic regions (**Fig. 3**). In contrast, 21 nt sRNA showed a peak density toward the center of gene bodies. The distributional patterns reported here resemble previously reported genic signatures of sRNA abundance in well-annotated genomes such as maize (Gent et al. 2013; Lunardon et al. 2016), *A. thaliana* (Dowen et al. 2012) and rice (Li et al. 2012). The relative decrease in transcript-associated 24 nt sRNA after stress exposure may suggests a loss of methylation in gene flanking regions (and possibly also in TE sequences within genes) and consequent gene expression upregulation. However, a quality genome reference assembly will be required to test this hypothesis. Together, the specific changes in sRNA profiles that we observed are in line with previous observations in stress-exposed plants, but our results indicate the stress-associated sRNA footprint is maintained transgenerationally for at least two unexposed generations after the stress treatment.

**Figure 3.**
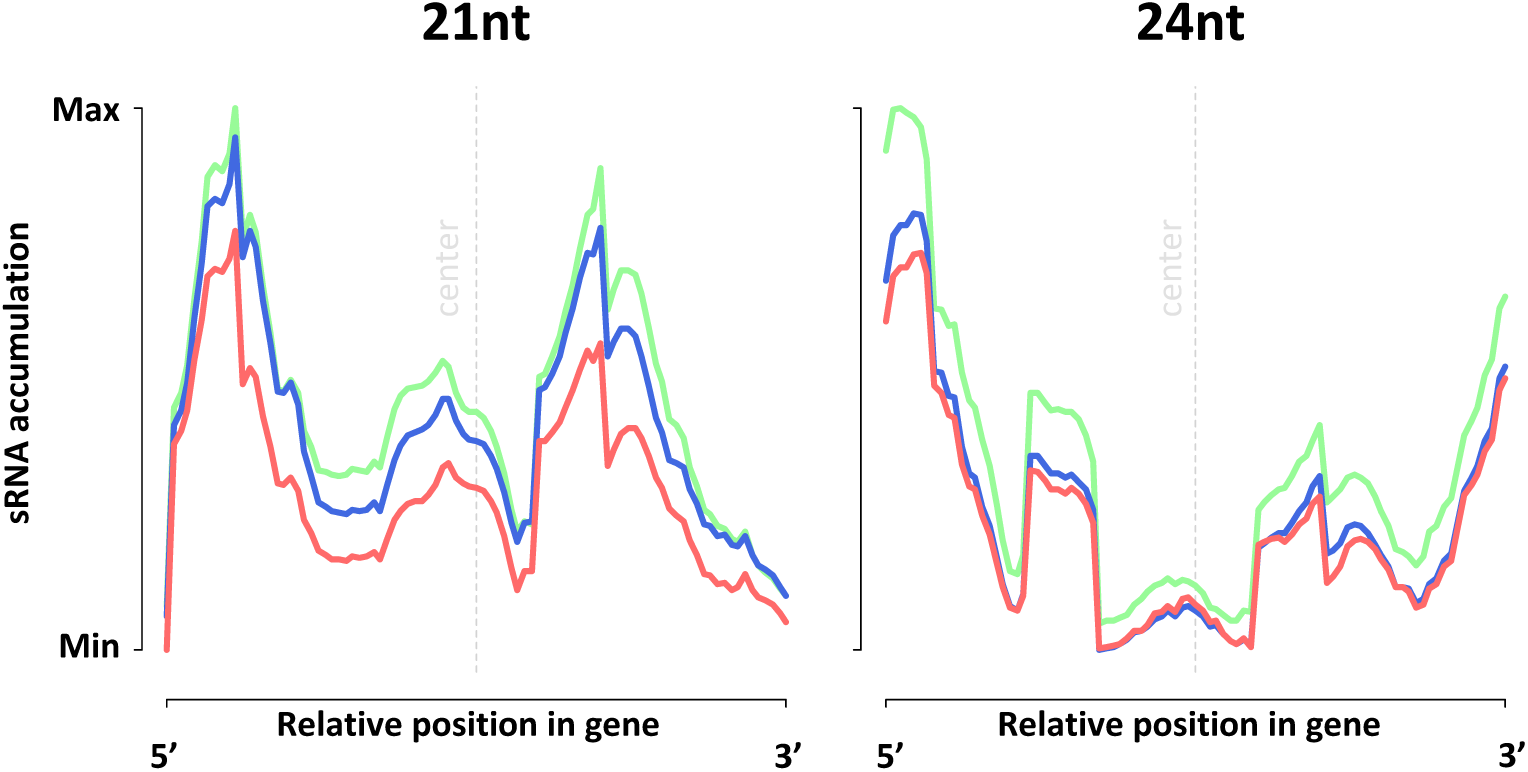
Spatial accumulation of 21 and 24 nt sRNA reads in gene-mapping transcripts. Lines represent density distributions of sRNA mapping location along the transcript. Each gene-mapping transcript was scaled to a length of 1000 bp and sRNA mapping positions (pooled replicates) inside each transcript were transformed accordingly. The counts for each transcript were afterwards collapsed into a single transcript model by calculating the averaged number of sRNA hits for each transcript coordinate across all length-normalized transcripts. Color code for treatment groups: control (green), drought (blue) and SA (red).

In order to identify specific genes that show different sRNA abundance comparing control and stress treatments, we performed differential analysis using DESeq2 (Love et al. 2014). We applied DESeq2 to different sRNA lengths: 21nt, 24nt sRNAs and for all length classes combined (18 to 30nt). After adjusting for multiple testing (FDR=0.10), our results showed virtually no significant sRNA enrichment or depletion at specific genes (Supplementary table: **S4**). We argue that the induced and transgenerationally inherited sRNA effects are subtle and may not be readily detectable using our approach that involved sRNA sequencing of individual plants, not pooled samples. We therefore focused, instead, on sets of genes that were either most depleted or enriched for sRNA in the stress groups and we tested if these gene sets were overrepresented for specific GO-terms. Our analysis revealed a strong overrepresentation for hundreds of GO categories, depending on grandparental treatment and sRNA length class (Supplementary text: **S5**). A large fraction of these GO-categories overlapped between the two stress treatments, suggesting a generalized response. Genes that were depleted or enriched for 21 nt sRNAs were significantly enriched for about 400-500 GO categories in both the control-drought comparison and in the control-SA comparison. For 24 nt sRNAs the downregulated genes were enriched for many more GO terms than the upregulated genes, suggesting a strong biological signal in the relaxation of 24 nt-based gene silencing after grandparental stress.

We searched the list of enriched GO terms for specific keywords that are associated with the grandparental stresses: “water” and “drought” for drought treatment, “salicylic” and “hormone” for SA treatment and “response to stress”, “abiotic stimulus” and “wounding” for stress treatments in general (see **Fig. 3**). For instance, the GO term 0006950 (“response to stress”) was specially enriched, pointing to an active stress memory. For all other key words, except “drought”, significantly enriched GO terms were found in both stress treatments, suggesting that these GO terms reflect a general stress response rather than a treatment-specific response. However, two SA-related GO categories (GO term 0009862: systemic acquired resistance, salicylic mediated signaling pathway; and GO term 0009914: hormone transport) were affected only in the SA set, indicating a more treatment-specific pattern.

In summary, it is well known that stress responses can be mediated by changes in sRNA-associated gene silencing. Our results suggest that this regulation may persist for several generations after stress. sRNA-based multi-generational inheritance of environmental stress has been previously demonstrated in some animal systems (e.g. Gapp et al. 2014; Rechavi et al. 2014) where underlying mechanisms of sRNA inheritance are at least partly different from plants. Although effects on gene expression remain to be evaluated, our study is to our knowledge the first demonstration in plants of modified sRNAs two generations removed from the stress trigger. Our results show no clear statistically significant effects on individual genes, which may be due to low sequencing depth of the libraries, or lack of sensitivity of our differential analysis. However, we were able to uncover a sRNA signal among genes involved in stress-related functions. This illustrates that an epigenetic signal travelled between generations preserving footprints of grandparental stress, and that this memory implicates genes that are known to be involved in stress responses. Although we did not explore the nature of the transgenerationally inherited epigenetic signal, this signal could be a stress-induced change in TE-associated DNA methylation, which in plants can be stably inherited and can trigger RNA-mediated gene expression changes in offspring (Wibowo et al. 2016).

## Acknowledgments and funding information

Keygene N.V., Wageningen, kindly provided *T. officinale* BAC sequences that we used for preliminary analysis of TE-mapping sRNAs. Seeds from the *T. officinale hemicyclum* lineage were kindly provided by Jan Kirschner from the CAS Institute of Botany, Pruhonice. This work was supported by the Netherlands Organisation for Scientific Research (grant numbers 864.10.008 and 884.10.003); and an ERC starting grant to O.R. (grant number 335624). sRNA data generated for this study are deposited in SRA, submission number SUB2306247.

## Supplementary text S1. Experimental methods

The experimental design is visualized in Figure S1. Apomictic dandelion plants from a single lineage (*T. officinale hemicyclum*) were used (Kirschner et al., 2016). G1 plants in the experiment derived through apomictic reproduction from a single G0 individual. Seedlings were distributed over three treatment groups: control, drought and salicylic acid. Plants within a group were randomized and placed in rows to ensure non-touching between the treatment groups. After four weeks of growth in a climate chamber (14 h light / 10 h dark, 18 °C / 14 °C, 60%) drought treatment started: during ten brief drought episodes over a period of four weeks, water was withheld until at least 80% of all individuals showed wilted leaves. At five weeks after seedling growth, SA treatment was applied once by spreading 0.5 ml of 10 mM SA solution over the surface of three leaves. The control group received no treatment. At eight weeks, plants were moved to a cold room for a 5-week vernalization period (continuous 4°C, 16 h light / 8 h dark), after which plants were placed on greenhouse benches and seeds were collected six weeks later. Using single-seed descent the subsequent two generations, G2 and G3, were grown in a common control environment under the same conditions as described for G1.

Leaf tissue was sampled from five weeks old G3 plants. Sixteen leaf discs of 8 mm in diameter were punched from one young and fully-developed leaf. Leaf discs were snap frozen in liquid nitrogen and stored in -80°C until usage. From liquid nitrogen-ground leaf tissue total RNA was extracted using 1 ml of Trizol (Ambion, Life technologies, The Netherlands) according to the manufacturer protocol but with an additional precipitation step with isopropanol and 3M sodium acetate (pH 5.2) and subsequent washing steps with ethanol. The final pellet was dissolved in 50 μ! DNase/RNase-free water. Quality was checked on agarose gel electrophoresis and concentration on a NanoDrop 1000 spectrophotometer. For RNA sequencing library preparation the New England Biolabs NEBNext multiplex small RNA kite7300 kit (Ornat, Rechovot, Israel) was used with an initial 1 μg of RNA according to the manufacturer protocol, using barcoded primers and using T4 ligase truncated for 3’ ligation. To select for sRNA with a length of 20-30nt, cuttings from a E-Gel EX 4% Agarose gel (Invitrogen, Life technologies, Israel) were taken between 140-150nt size fraction (accounting for two adapter sequences of 60nt each). Bands were cleaned using MiniElute Gel Extraction kit (QIAGen, Eldan, Israel). After clean-up and a final Bioanalyzer quality check (Agilent Technologies, Eldan,Israel) using Agilent High Sensitivity DNA Kit, the 12 samples were pooled into a single sequencing library which was sequenced on two Illumina Hiseq2500 lanes (50bp, single-end).

---

**Figure S1.**
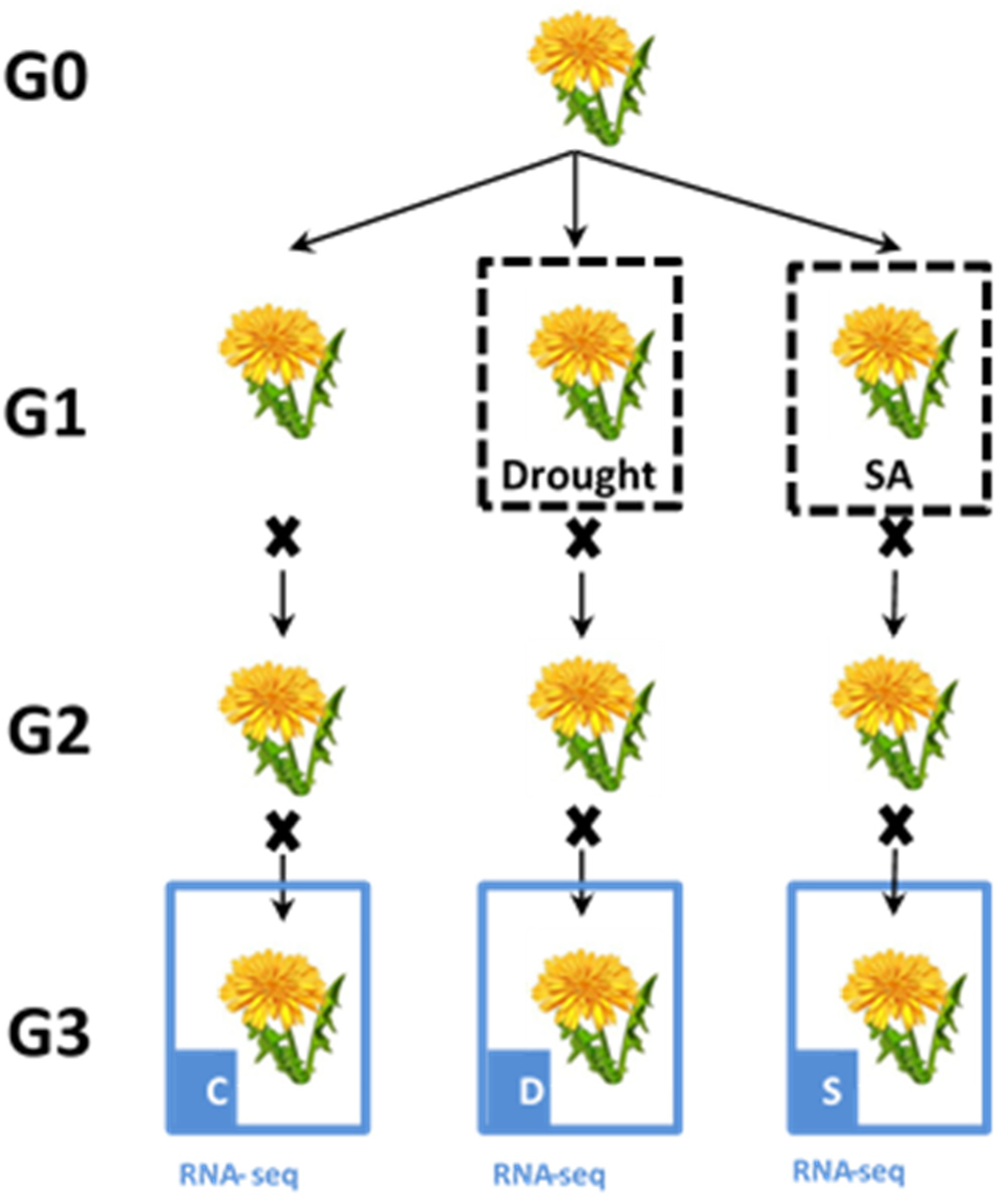
Apomictic Dandelion lines generated for the current study. Generations 1 to 3 are marked by G1, G2 and G3. Each plant cartoon accounts for 4 replicates, except for the single initial common ancestor individual (G0). Lines subjected to sRNA sequencing are indicated with a blue box. Treatment sets: control (C), drought (D); salicylic acid (S).

## Supplementary text S2. Data processing and mapping

FastQC v0.11.3 software (www.bioinformatics.babraham.ac.uk/projects/fastqc/) was used for preliminary quality check. Reads were processed using adapter trimming Cutadapt (Martin, 2011), filtered for quality (Phred score>33), and length (18nt to 30nt). Read counts per sRNA showed overall good correlations between replicate samples within experimental groups (Fig. S2). Reads were aligned with BWA (Li and Durbin, 2009; perfect matches) to:

1. A *Taraxacum officinale* reference transcriptome (Ferreira de Carvalho et al., 2016b). The transcriptome was assembled *de novo* from RNAseq data, which resulted in a total of 123,232 transcripts of which 39,685 transcripts were annotated to TAIR genes (BLASTn, e-value< 1e-05). The transcriptome is available in the Dryad repository under identifier number doi:10.5061/dryad.6p2n6.
2. A *Taraxacum officinale* database of transposable element sequences. This TE database was based on *de novo* clustering of genomic sequencing reads of the apomict *T. officinale* ‘macranthoides accession 12’ using RepeatExplorer, as presented in Ferreira de Carvalho et al. (2016a) and available in the Dryad repository under identifier number doi:10.5061/dryad.f2j00. From the list of clusters assembled by RepeatExplorer, the most abundant clusters (i.e. containing at least 0.20% of the inputs reads) were selected. Thus, a total of 120 clusters were used for sRNA mapping and represent highly repetitive annotated TE sequences present in the genome of *T. officinale.*

**Figure S2.**
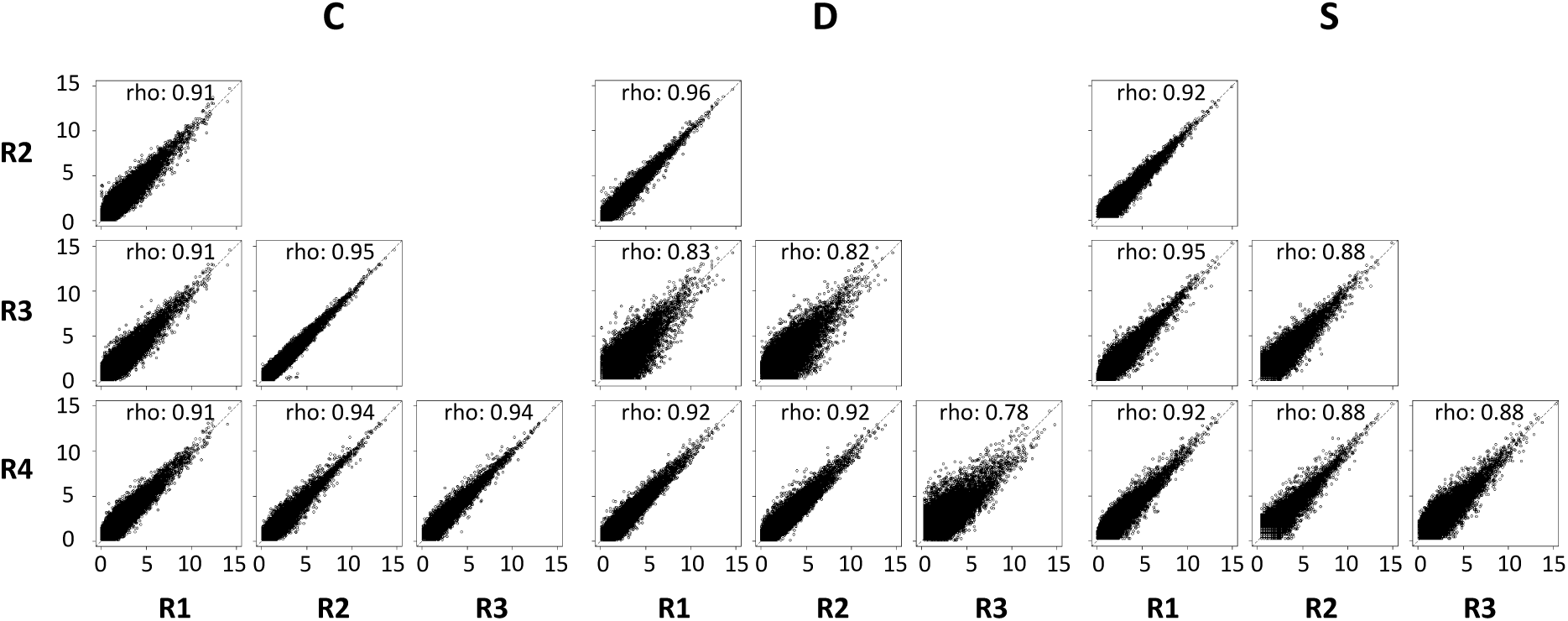
Correlation between sRNA read counts among replicate libraries (R1, R2, R3 and R4) in each treatment group (C, D and S). Axes in each plot represent the log2(RPM+1). Rho: Pearson correlation coefficient based on read count of sRNAs that are shared between the two libraries.

## Supplementary text S3. Bootstrap procedure

The bootstrap procedure to determine significance of global shifts in sRNA composition among treatment sets compared a pooled control with a pooled treatment sample (drought or salicylic acid); in both groups the 4 replicate plants were pooled in silico into a single sample. Generically, the bootstrap procedure for hypothesis testing implemented considers two observed independent samples:

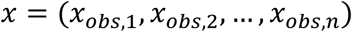

and

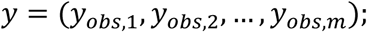

with *n* and *m* the respective sizes, and for which no assumptions of normality must be made. From these observed samples the mean difference

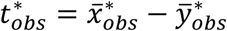

can be calculated, for which the null hypothesis

*H_0_: both samples come from the same population* is tested against the alternative

*H_1_. both samples are NOT from the same population.*

The method encompasses 5 steps:

1. Merging of the two observed samples into one sample of (*n +* m)observations
2. A bootstrap of (*n +* m)observations with replacement is drawn from the merged sample
3. The mean *x^̅*^* for the first *n* observations is calculated and then the mean *y^̅*^* for the remaining *m* observations, afterward which the test statistic *t^*^* = *x^̅*^* — *y^̅*^* is calculated
4. Steps 2 and 3 must be repeats B (e.g. 4000) times to obtain B values of the test statistic
5. A p-value is then estimated according to:

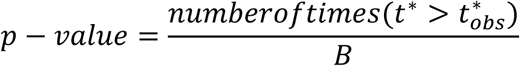

If *p* — *value* < *α*, being *a*the significance level (e.g.: 0.05) then *H_0_is* rejected, and retained otherwise.

## Supplementary text S4. DESeq2 results

**Table S4.** Per-transcript analysis of differential sRNA abundance. DESeq2 results for: all consensus sRNA (sRNA shared by all replicates from at least one of the treatments), TE-mapping sRNA and gene-mapping sRNA. Up- or downregulated sRNAs are significant after multiple testing correction in DESeq2 at FDR = 0.1 threshold.

| Group | Comparison | sRNA length (nt) | #instances tested | #Up-regulated | #Down-regulated |
| --- | --- | --- | --- | --- | --- |
| Consensus set | D/C | All | 569268 | 3 | 0 |
|  |  | 21 | 102759 | 2 | 0 |
|  |  | 24 | 235625 | 0 | 0 |
|  | S/C | All | 545459 | 15 | 12 |
|  |  | 21 | 99072 | 5 | 8 |
|  |  | 24 | 227383 | 0 | 6 |
| TEs | D/C | All | 98 | 0 | 0 |
|  |  | 21 | 98 | 0 | 0 |
|  |  | 24 | 98 | 0 | 0 |
|  | S/C | All | 98 | 0 | 0 |
|  |  | 21 | 98 | 0 | 0 |
|  |  | 24 | 98 | 0 | 0 |
| Gene-mapping transcripts | D/C | All | 35880 | 0 | 5 |
|  |  | 21 | 27153 | 0 | 0 |
|  |  | 24 | 27730 | 0 | 1 |
|  | S/C | All | 36202 | 0 | 0 |
|  |  | 21 | 27975 | 0 | 0 |
|  |  | 24 | 28683 | 0 | 0 |

## Supplementary text S5. Gene Ontology analysis

Gene Ontology (GO) analysis was performed with the R package topGO (Alexa and Rahnenfuhrer, 2004) and annotations from the Arabidopsis genome (Carlson *GO.db* v.3.1.2). To control for potential biases due to the use of Arabidopsis as a reference in the enrichment analysis, a baseline enrichment level was determined via random subsampling from the dandelion transcripts. This subsampling was repeated 1000 times picking a number of random transcripts equal to the top 5% of the respective treatment-control set (control versus drought and control versus SA). We considered the enrichment to be significant if the FDR-adjusted p-values from the topGO enrichment analyses exceeded the 95% confidence interval of the bootstrapped subsamples.

**Figure S5.**
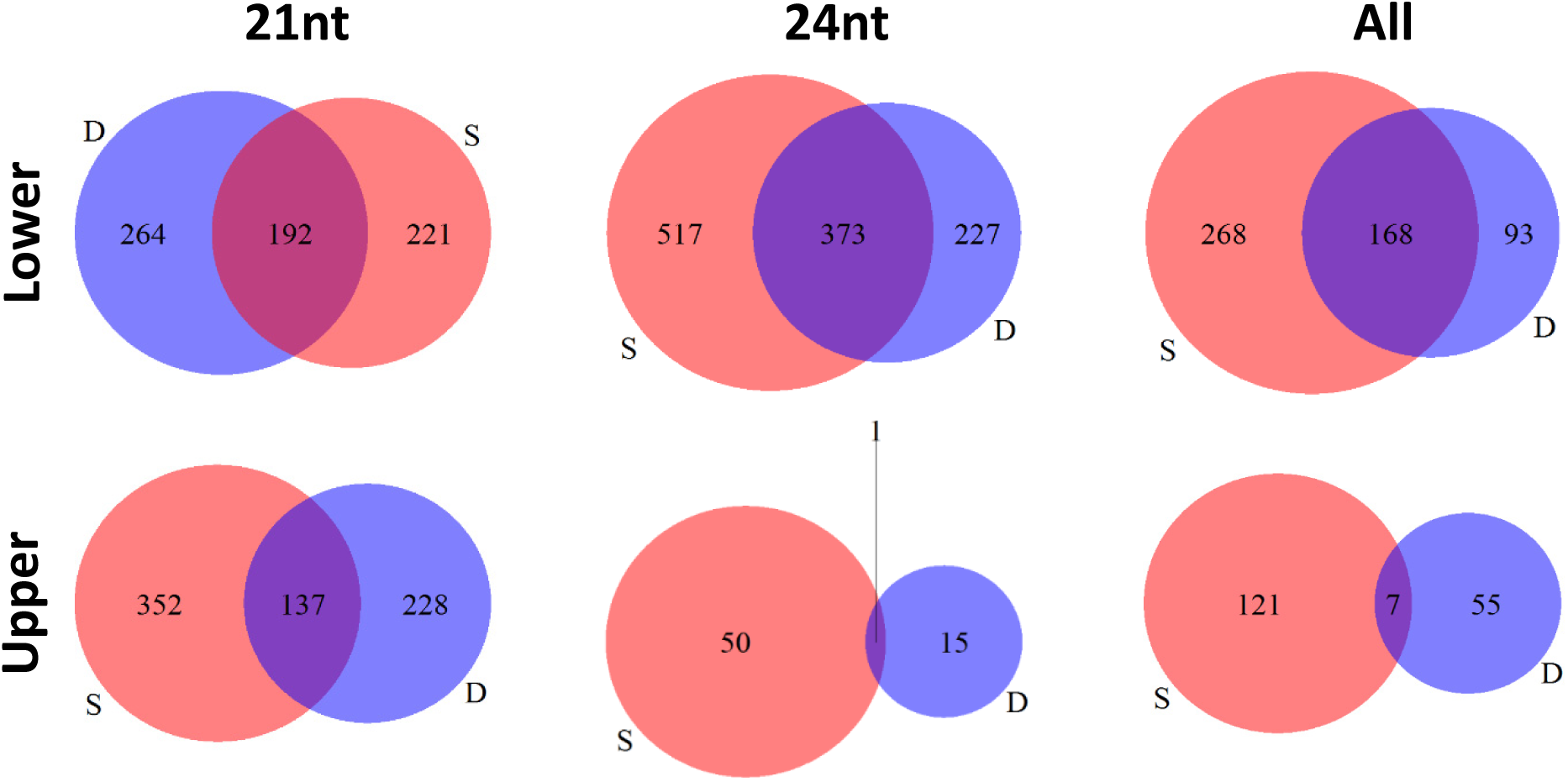
Overlap between GO terms that are significantly enriched in the list of most downregulated genes (lower: 5% of transcript with strongest reduction in mapped sRNAs due to grandparental stress compared to control treatment) and in the list of most upregulated genes (upper: 5% of transcripts with strongest increase in mapped sRNA due to grandparental stress treatment). Numbers for significantly enriched GO terms are shown as Venn diagrams for effects of grandparental drought and grandparental SA treatment for 21nt and 24nt sRNAs and for all length classes between 18 and 30nt.

**Table S5.** Number of genes for each of the 11 terms selected in the GO analysis.

| GO id | Term | #Annotated genes | #Significant genes |  |  |  |
| --- | --- | --- | --- | --- | --- | --- |
|  |  |  | CvsD |  | CvsS |  |
|  |  |  | Lower 5% | Upper 5% | Lower 5% | Upper 5% |
| GO:0009414 | response to water deprivation | 419 | 57 | 28 | 52 | 31 |
| GO:0009751 | response to salicylic acid | 471 | 53 | 25 | 55 | 34 |
| GO:0009862 | systemic acquired resistance, salicylic ... | 251 | 24 | 13 | 33 | 21 |
| GO:0009725 | response to hormone | 1601 | 145 | 100 | 148 | 120 |
| GO:0009755 | hormone-mediated signaling pathway | 773 | 82 | 58 | 88 | 66 |
| GO:0009914 | hormone transport | 107 | 14 | 13 | 13 | 19 |
| GO:0006950 | response to stress | 4030 | 406 | 280 | 398 | 286 |
| GO:0080134 | regulation of response to stress | 569 | 77 | 34 | 77 | 46 |
| GO:0080136 | priming of cellular response to stress | 2 | 1 | 0 | 1 | 1 |
| GO:0009628 | response to abiotic stimulus | 2796 | 307 | 208 | 326 | 197 |
| GO:0009611 | response to wounding | 335 | 57 | 25 | 44 | 38 |

